# Rethinking Representation Complexity in Drug–Target Prediction via Supervised Vector Quantization

**DOI:** 10.1101/2025.06.13.659523

**Authors:** Jiandong Chen, Yao-zhong Zhang, Lu Lu, Meixi Wu, Zhiang Chen, Seiya Imoto, Chen Li

**Affiliations:** Department of Human Genetics and Women’s Hospital, Zhejiang University School of Medicine, 866 Yuhangtang Road, Hangzhou 310058, China; Division of Health Medical Intelligence, Human Genome Center, The Institute of Medical Science, The University of Tokyo, Shirokanedai 4-6-1, Minato-ku, Tokyo 108-8639, Japan; College of Computer Science, Jiangxi University of Chinese Medicine, 1688 Meilin Avenue, Nanchang 330004, China; Department of Computational Biology and Medical Sciences, Graduate School of Frontier Sciences, The University of Tokyo, Shirokanedai 4-6-1, Minato-ku, Tokyo 108-8639, Japan

**Author notes:** Corresponding author: Yao-zhong Zhang, Seiya Imoto and Chen Li.

**Keywords:** Drug-target interaction, Supervised vector quantization, Deep learning, Protein language model

## Abstract

Accurate prediction of drug–target interactions (DTIs) is crucial for computational drug discovery. Although pretrained language models offer richer molecular and protein representations, their increasing complexity does not always lead to better predictive performance. In many cases, the inclusion of redundant or irrelevant features may obscure biologically relevant patterns. In this study, we systematically evaluate the contribution of complex features in DTI prediction and demonstrate that only a portion of these features is truly informative. Based on this insight, we propose a Vector Quantization (VQ)-based module that functions as a plug-and-play feature selection layer within deep learning architectures. When combined with a simple fully connected classifier, this supervised VQ (SVQ) framework not only surpasses recent state-of-the-art DTI methods in performance, but also enhances interpretability through the learning of discriminative codewords. This work highlights the importance of input feature selection in deep learning and offers a new perspective for constructing robust and interpretable DTI prediction models.

## 1 Introduction

In the complex and resource-intensive process of drug discovery, one of the most costly and time-consuming steps is the large-scale screening of small molecules against protein targets. To reduce these costs and improve efficiency, computational approaches have become valuable alternatives to experimental screening. These methods generally fall into two categories: molecular docking (MD) and machine learning-based prediction models [1]. MD approach simulates the physical interactions between molecules and target proteins in silico, offering improved throughput compared to experimental assays. However, it heavily depends on the availability of protein cocrystal structures [2], and individual calculations are still required for each molecule–target pair, both of which limit its scalability for large-scale screening. In contrast, machine learning-based methods employ fixed-size models trained on existing drug-target interaction (DTI) data, significantly enhancing computational efficiency. Once trained, these models can rapidly predict potential interactions across large chemical libraries without exhaustive pairwise calculations, making them a more attractive alternative that has gained rapid development in recent years.

To advance machine learning-based DTI prediction, a variety of computational strategies have been developed. Among them, similarity-based approaches offer an intuitive framework by assuming that structurally similar drugs tend to share similar targets. These methods typically compute similarities based on chemical structures and protein sequences, and incorporate them into frameworks such as nearest-neighbor algorithms, network-based inference methods, and Support Vector Machines (SVM) [3]. These approaches often yield strong predictive performance, achieving high area under the precision–recall curve (AUPR) and area under the receiver operating characteristic curve (AUROC) scores. However, their reliance on predefined similarity metrics limits their ability to generalize to chemically dissimilar or novel compounds. Meanwhile, feature-based methods provide a similarity-independent approach by encoding drugs and targets into fixed-length feature vector representations. This approach enables the capture of broader biochemical and contextual information beyond simple drug–drug and target–target similarities, allowing for more flexible and potentially novel DTI predictions.

Building on the strengths of feature-based strategies, the representations of drugs and proteins have progressed significantly, evolving from manually curated descriptors to more expressive and data-driven encodings. For example, Zhang *et al*. extracted database descriptors (e.g. hydrophobicity index) as features for drugs and proteins [4]. Later, Wang *et al*. encoded drugs using molecular fingerprints and represented target proteins with a method called the position-specific scoring matrix (PSSM), providing more comprehensive representations for both drug compounds and proteins [5]. With the development of deep learning, representation learning has become increasingly prominent. Lee *et al*. proposed a convolutional neural network (CNN)-based framework that integrates Morgan fingerprints of drugs with protein embeddings generated from randomly encoded amino acid sequences [6]. More recently, DrugBAN further combined CNNs and graph convolutional networks (GCNs) to model protein sequences and the local structures of drug molecules, achieving better predictive performance than earlier models [7].

In recent years, the emergence of sequence-based pretrained models has facilitated more advanced representation techniques for both drugs and proteins. Models such as MoLFormer [8] and ChemBERTa [9] for compounds, and ProtBERT [10] and ESM-2 [11] for proteins, typically developed for general representating purpose, are pretrained on large-scale molecular or amino acid sequences via self-supervised masked token prediction. In this way, these models can effectively capture not only local structural or sequential features but also long-range dependencies, thereby enriching the encoded representations. Several recent studies have incorporated such pretrained representations into DTI prediction frameworks. For instance, ConPLex utilizes ProtBERT-derived protein embeddings and introduces a protein-anchored contrastive learning scheme, achieving competitive performance [12]. Koyama *et al*. further proposed ChemGLaM, which takes advantage of the rich representations provided by MoLFormer and ESM-2 for drug and protein, respectively, and fine-tunes them using a cross-attention interaction block [13].

In conclusion, with the evolution of representing techniques, drug and protein embeddings have become increasingly comprehensive and dense in features. For years, the notion that “more is better” has been widespread, suggesting that incorporating more features would inevitably lead to better model performance. However, this assumption does not necessarily hold true in all cases. Typically, in specific tasks with a limited sample size, such as DTI prediction that involves complex biomedical data, a large feature set may increase the risk of model overfitting. Yet, as systematically identifying the most informative features for these tasks remains a challenge, researchers conventionally feed the model with all available features and rely on the model to discriminate feature relevance. This raises critical questions: how many of these pretrained, general-purpose features are truly necessary? And if a substantial portion is irrelevant or redundant, would it be beneficial to eliminate them prior to model input?

Motivated by these questions, we conducted a systematic investigation into the contributions of different complex representation features under the context of DTI prediction. Our study examined the effect of prior feature selection on model performance and revealed substantial improvements when irrelevant or redundant features were filtered out. Encouraged by these findings, we proposed a plug-and-play module that facilitates end-to-end, supervised “soft” feature selection within deep learning workflows. This module exploits the inherent properties of Vector Quantization (VQ) [14] to compress complex representations and suppress irrelevant noise. When combined with a simple fully connected layer, we developed a supervised VQ (SVQ) framework that not only surpasses recent state-of-the-art DTI methods in performance but also enhances interpretability via discriminative codeword learning. The main contributions of this study are summarized as follows:

1. We demonstrate that not all features within complex representations are critical for effective model learning, and that selectively filtering of low-value features can led to significant performance improvements.
2. We propose a VQ-based plugin module approximating feature selection in an end-to-end supervised learning framework. Integrated with a simple fully connected layer, this SVQ framework achieves state-of-the-art performance while improving interpretability.
3. We demonstrate the advanced interpretability of the SVQ framework by showing that, through supervised learning, it encodes typical DTI interaction patterns directly within its codeword usage distribution without any additional decoding process. Further analysis revealed that interaction-relevant information is stored primarily in the co-occurrence patterns of codewords rather than in their explicit semantic contents, suggesting a broader paradigm in deep learning: emphasizing co-occurrence patterns over precise embeddings can lead to efficient and interpretable models.

## 2 Results

### 2.1 Not all features from pretrained language models are essential for DTI prediction

Pretrained language models often need to balance general-purpose transferability with task-specific discriminability. In DTI prediction, these models often generate high-dimensional representations with numerous features, many of which are irrelevant to the task, introducing redundancy that can obscure informative signals and hinder model performance. To address this issue, we explore feature reduction to enhance the discriminative capacity of the representations. In particular, we leverage Random Forest (RF), which intrinsically estimates feature importance and facilitates the selection of informative features, making it a suitable tool for this investigation. Specifically, we applied an RF classifier to three widely used datasets—BioSNAP, BindingDB, and DAVIS—using 512 drug features and 1,152 target features extracted from pretrained language models for each drug-target interaction pair.

In our experiments, compared to state-of-the-art methods such as ConPLex, EnzPred-CPI [15], MolTrans [16], and DeepConv-DTI [17], the RF classifier demonstrated strong competitiveness in classifying DTI (Table 1). Notably, it achieved superior performance in terms of AUPR, demonstrating the effectiveness of feature selection in improving model performance.

**Table 1:** Comparison of AUPR scores between RF and recent DTI prediction methods.

| Model | BIOSNAP | BindingDB | DAVIS |
| --- | --- | --- | --- |
| ConPLex | $0.897 \pm 0.001$ | $0.628 \pm 0.012$ | $0.458 \pm 0.016$ |
| EnzPred-CPI | $0.866 \pm 0.003$ | $0.602 \pm 0.006$ | $0.277 \pm 0.009$ |
| MolTrans | $0.885 \pm 0.005$ | $0.598 \pm 0.013$ | $0.335 \pm 0.017$ |
| DeepConv-DTI | $0.889 \pm 0.005$ | $0.611 \pm 0.015$ | $0.299 \pm 0.039$ |
| **RF** | **$0.915 \pm 0.001$** | **$0.669 \pm 0.002$** | **$0.486 \pm 0.005$** |

**Table 2:** Model performance comparison applying SVQ.

| Model | BIOSNAP | BIOSNAP<br>unseen drugs | BIOSNAP<br>unseen targets | BindingDB | DAVIS |
| --- | --- | --- | --- | --- | --- |
| MLP | 0.924 ± 0.001 | 0.902 ± 0.001 | 0.844 ± 0.001 | 0.657 ± 0.001 | 0.447 ± 0.007 |
| Boruta-MLP | 0.926 ± 0.000 | 0.907 ± 0.002 | 0.841 ± 0.005 | 0.666 ± 0.004 | 0.448 ± 0.010 |
| **SVQ-MLP** | **0.928 ± 0.000** | **0.900 ± 0.002** | **0.835 ± 0.003** | **0.668 ± 0.006** | **0.456 ± 0.011** |

To identify the key features relevant to DTI, we employed the Boruta feature selection algorithm along with the RF classifier [18]. Boruta works by introducing randomly shuffled “shadow” features together with their original counterparts during RF classifier training, and then discarding any features whose contribution is lower than that of the shadow features. The RF classifier uses Gini impurity to assess feature importance. After 10 iterations of Boruta permutation, fewer than 30% of the original features were retained (26.2% for BioSNAP, 15.2% for BindingDB and 4.3% for DAVIS as shown in Figure 1). This significant reduction indicates that the majority of features do not contribute meaningfully to the DTI task, and only a small subset of features are truly informative for DTI prediction.

**Figure 1:**
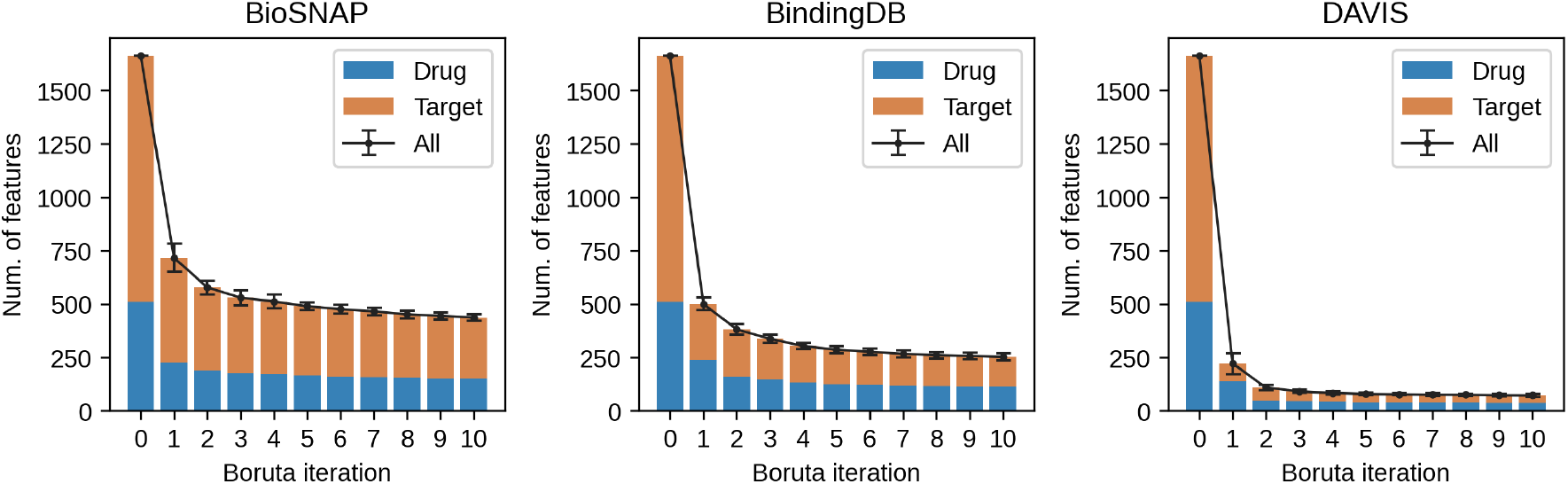
Number of features retained after each Boruta iteration for DTI prediction.

### 2.2 Simple feature selection enhances feature quality for DTI prediction

While these selected features show higher importance within the RF classifier, it remains unclear whether this corresponds to genuinely higher feature quality. To verify this, we employed a linear probing approach [19], in which a simple multilayer perceptron (MLP) was appended to the selected feature inputs to evaluate their impact on downstream performance. As a result, the MLP classifier’s performance improved significantly when trained with these features in both BioSNAP and BindingDB (Figure 2). In contrast, blocking the classifier from accessing these features led to a notable performance drop on both datasets.

**Figure 2:**
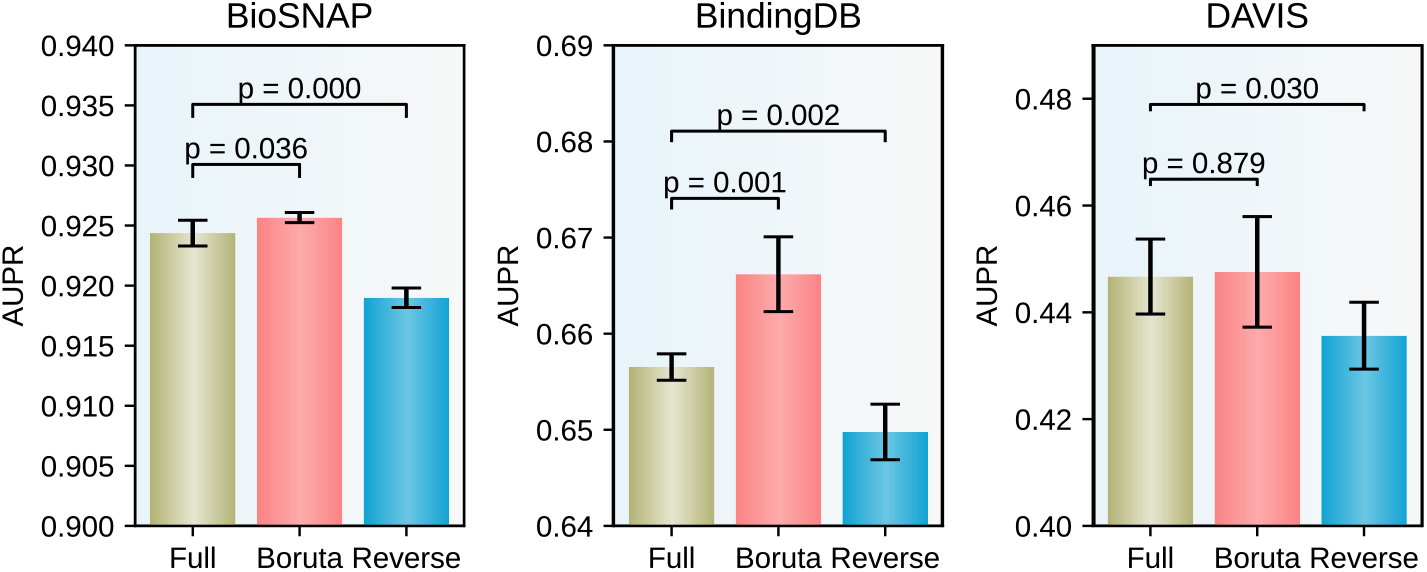
Performance comparison of linear probing using different feature inputs. All: full feature set; Boruta: features retained after 10 Boruta iteration; Reverse: features discarded by Boruta. Statistical significance tested by Student’s t-test.

In the DAVIS dataset, using Boruta-selected features did not lead to a substantial performance gain. Nevertheless, excluding these features caused a significant decline in performance. This indicates their essential role, even if their individual contribution appears less prominent. It is also worth noting that the effectiveness of RF-based feature selection depends on the size and diversity of the training data. The relatiely small number of drugs and proteins in DAVIS may limit the model’s ability to adequately identify informative features.

### 2.3 Application of SVQ significantly enhances model performance in DTI prediction

While the RF-based feature selection strategy has demonstrated effectiveness in improving DTI prediction, it also presents inherent limitations. First, such conventional machine learning workflow is difficult to dynamically incorporate into end-to-end deep learning architectures. Second, it tends to underestimate the value of distributed representations, where information is encoded not in isolated dimensions but through interactions among multiple features with synergistic or superposition effects. As a result, a tree-based feature selection method is inherently not well-suited for this pattern.

A promising alternative is VQ on spliced representation, which offers both feature selection in an oblique way and deep learning framework compatibility. It works by dynamically mapping spliced subspaces of feature sets into a limited set of learnable discrete vectors, also known as the codewords of VQ codebook, with each serving as a prototype and a representative anchor that ultimately summarizes the underlying subspace. This adds a discrete constraint to the feature distribution, reducing representation redundancy. Furthermore, the aggregation of quantized subspaces eventually gives rise to a higher-level prototype space, enabling the maintenance of the underlying depth of biomedical knowledge while achieving the feature selection goal.

Based on these considerations, we propose an SVQ framework, in which a plug-and-play VQ module with splicing implementation replaces the Boruta-dependent feature selection in the baseline MLP model (Figure 3). By learning a representative discrete representation, the VQ module emphasizes the most contributive features, offering a structured and interpretable approach to capturing critical interactions within the DTI space. In our benchmarks, incorporating the SVQ module substantially recovered the performance of the baseline MLP model, which otherwise showed notable degradation without input feature selection. Moreover, SVQ achieved a modest yet consistent performance improvement over the Boruta-enhanced model, suggesting its effectiveness in compressing high-dimensional features while retaining discriminative capacity.

**Figure 3:**
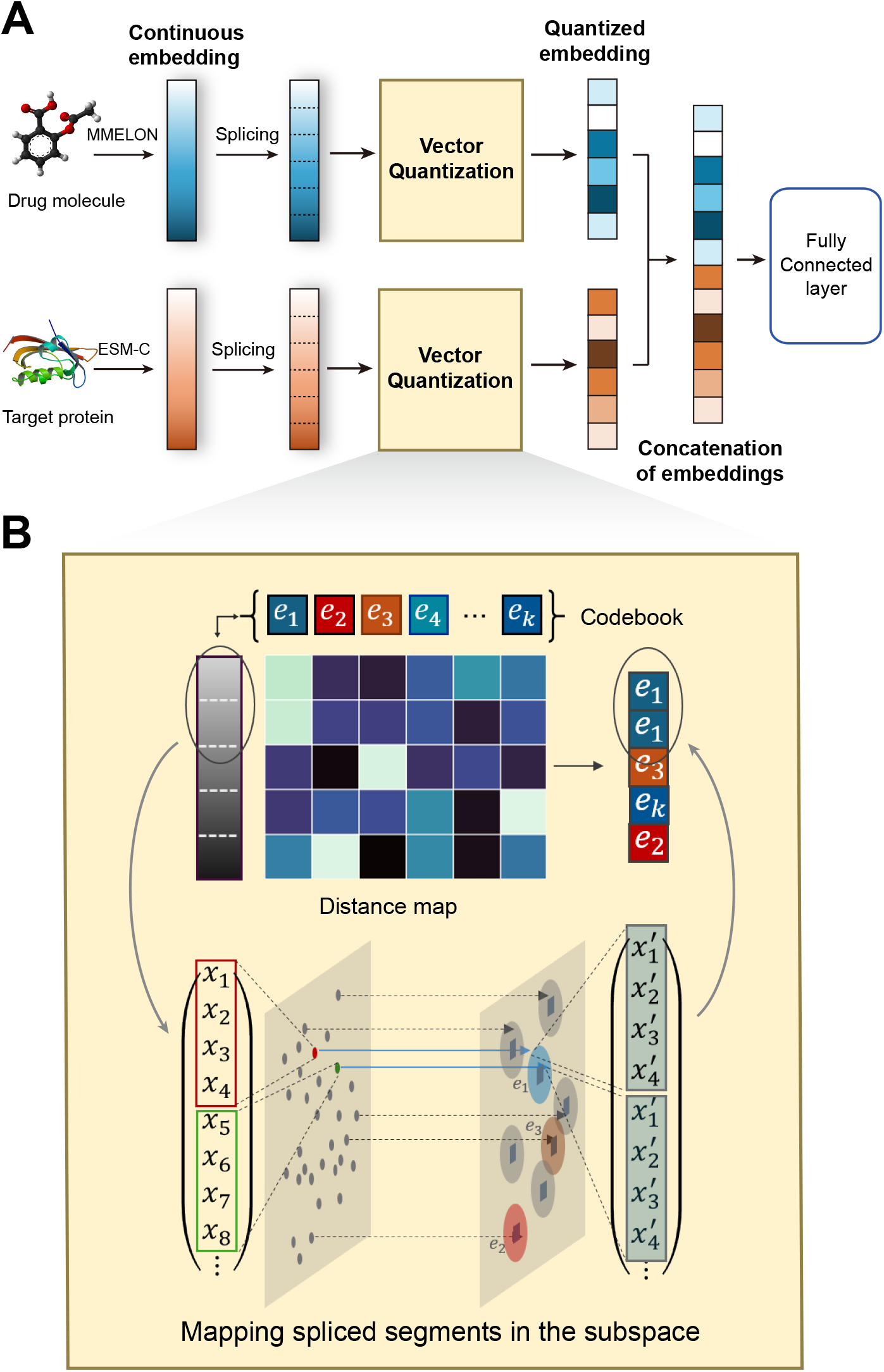
Schematic overview of the SVQ framework for DTI prediction. A. SVQ framework integrates a spliced VQ module into the baseline MLP architecture, enabling structured compression and selection of informative features for improved performance and interpretability. MMELON: Multi-view Molecular Embedding with Late Fusion model; ESM-C: Evolutionary Scale Modeling Cambrian. B. In the subspace of spliced embedding, each continuous segment is projected onto a nearest codeword from a discrete codebook, effectively quantizing the representation.

Notably, the SVQ-integrated model did not exhibit superior performance under zero-shot settings. When presented with previously unseen drugs and targets, SVQ’s advantage diminished, and both the plain and Boruta-enhanced models performed better. This outcome may reflect a trade-off introduced by the VQ module: while it effectively constraints known representations to reduce complexity and redundancy, it also and enhance interpretability, such constraints may reduce the model’s flexibility in handling out-of-distribution inputs. Nevertheless, SVQ consistently outperformed several recent state-of-the-art methods and offers an additional benefit of interpretability that Boruta could not offer, which will be further discussed in the subsequent section.

### 2.4 Comparative evaluation of SVQ and recent deep learning-based DTI methods

We compared SVQ with several recently proposed neural network-based methods that have demonstrated strong performance on DTI prediction tasks. As shown in Table 3, SVQ consistently exhibited moderate to strong performance advantages across the majority of datasets.

**Table 3:** Performance comparisons with recent state-of-the-art models.

| Model | BIOSNAP | BIOSNAP<br>unseen drugs | BIOSNAP<br>unseen targets | BindingDB | DAVIS |
| --- | --- | --- | --- | --- | --- |
| ConPLex | $0.897 \pm 0.001$ | $0.874 \pm 0.002$ | $0.842 \pm 0.006$ | $0.628 \pm 0.012$ | $0.458 \pm 0.016$ |
| EnzPred-CPI | $0.866 \pm 0.003$ | $0.844 \pm 0.005$ | $0.795 \pm 0.004$ | $0.602 \pm 0.006$ | $0.277 \pm 0.009$ |
| MolTrans | $0.885 \pm 0.005$ | $0.863 \pm 0.005$ | $0.668 \pm 0.045$ | $0.598 \pm 0.013$ | $0.335 \pm 0.017$ |
| DeepConv-DTI | $0.889 \pm 0.005$ | $0.847 \pm 0.009$ | $0.766 \pm 0.022$ | $0.611 \pm 0.015$ | $0.299 \pm 0.039$ |
| **SVQ** | **$0.928 \pm 0.000$** | **$0.900 \pm 0.002$** | **$0.835 \pm 0.003$** | **$0.668 \pm 0.006$** | **$0.456 \pm 0.011$** |

In the BIOSNAP dataset, SVQ achieved an AUPR of 0.928 ± 0.000, surpassing the next-best method, ConPLex, by 3.5%. A similar trend was observed in BindingDB, where SVQ reached an AUPR of 0.668 ± 0.006, outperforming ConPLex by 6.4%. These results highlight the competitiveness of SVQ particularly in improving predictive accuracy. In the DAVIS dataset, while SVQ did not exceed ConPLex (0.456 vs. 0.458), its performance remained closely comparable, demonstrating its robustness across datasets with Mapping spliced segments in the subspace different characteristics.

In zero-shot setting, SVQ showed notable effectiveness in the unseen drug scenario, achieving the highest AUPR (0.900 ± 0.002) among all compared models. In the unseen target scenario, SVQ slightly under-performed relative to ConPLex (0.835 vs. 0.842), yet still outperformed the remaining baselines, further affirming its strong generalization capability.

### 2.5 Domain Knowledge Emerges in SVQ Representations of Drug–Target Pairs

Given the strong performance of the SVQ framework, we speculate that the VQ module may encode valuable information relevant to drug-target interactions. If this is the case, the codeword usage associated with a given drug should correlate with the characteristics of its interacting proteins.

Protein domains are fundamental structural and functional units and are often the primary targets of drugs molecules. Many therapeutic strategies have focused on specific domains. For instance, the kinase inhibitor imatinib targets the ATP-binding domain of the Abl kinase [20]. Protein language models, which are trained on large-scale protein sequence data, are expected to capture such domain-related information. Recent studies using ESM model series have demonstrated their ability to infer protein domains without relying on prior annotations [21, 22], providing support for this assumption. As shown in Figure 4, visualization of embeddings generated for representative proteins revealed distinct clusters in the embedding space, with each cluster corresponding to a specific domain. This indicates that domain-level structure is indeed reflected in the learned representations of these models.

**Figure 4:**
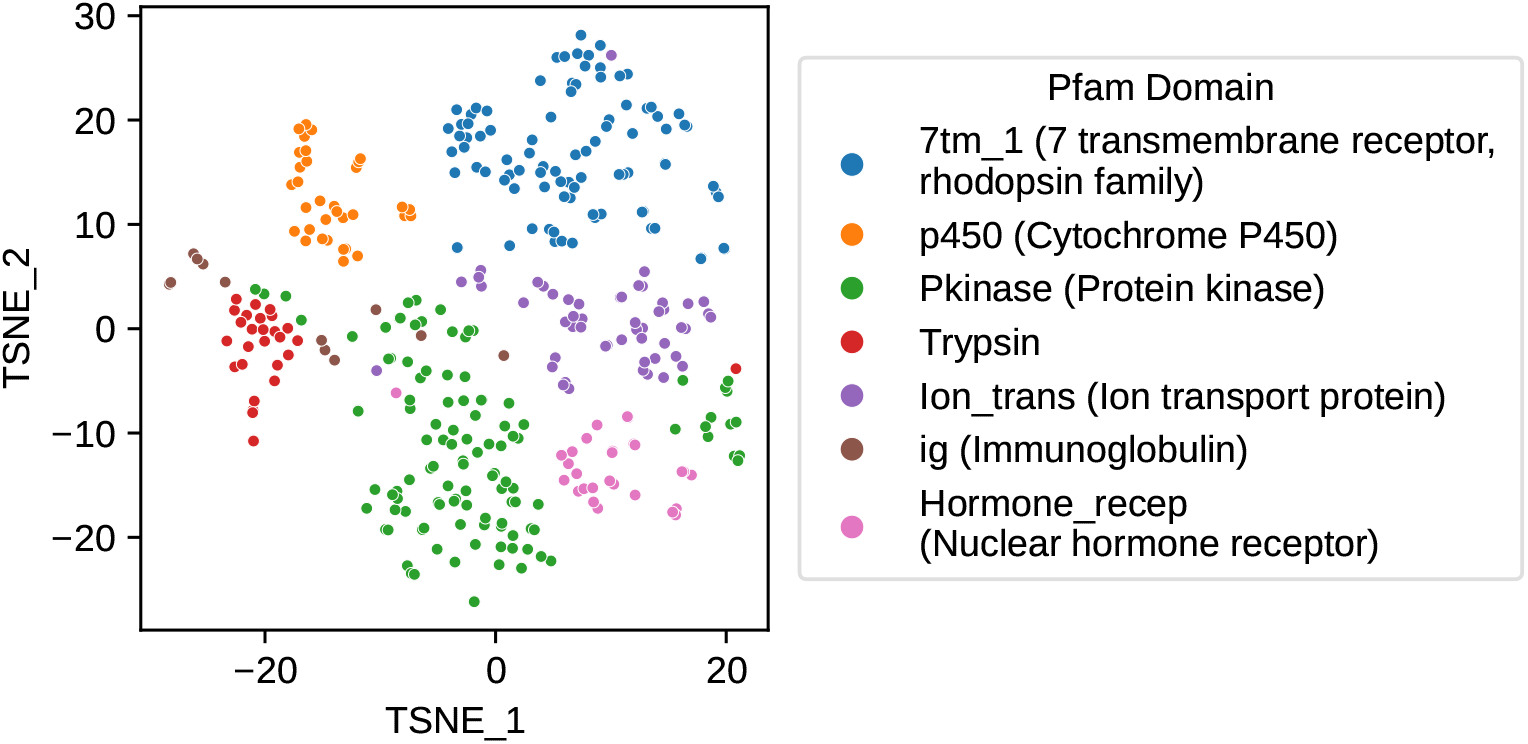
t-SNE projection of protein embeddings associated with unique Pfam domains. Proteins sharing the same domain tend to form clusters in the embedding space.

Motivated by these observations, we investigated whether the learned codebook exhibits domain-specific patterns in codeword usage. To this end, we analyzed the distribution of codeword usage among drugs, stratified by their interaction with specific protein domains. Each drug embedding was treated as a document composed of discrete codewords, and we applied a term frequency–inverse document frequency (TF-IDF) transformation to quantify the relative importance of each codeword across the drug set. To compare the relevance of codewords between interacting and non-interacting drugs, we defined the TF-IDF ratio score as

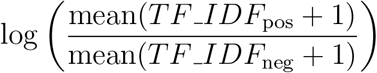

where *TF IDF*_pos_ and *TF IDF*_neg_ refer to the TF-IDF scores from drugs interacting with or not interacting with the domain. A higher ratio suggests that specific codewords are preferentially used by drugs targeting the domain, indicating that SVQ captures domain-relevant representations.

As a result, the learned codebook exhibited distinct usage patterns across different types of protein domains. As shown in Figure 5, Pfam domains [23] belonging to the same functional category tended to cluster together based on their associated codeword usage profiles. For instance, Module A highlighted a set of 15 codewords consistently underutilized by drugs targeting immunoglobulin-interacting domains, such as *V-set, ig, ig 3, I-set*, and *Ig 2* ). In contrast, Module B, comprising 16 codewords, was associated with ligand-gated ion channels, including *ANF receptor, Lig chan, Lig chan-Glu bd*, and other neurotransmitter-gated channels, which showed high usage frequencies in interacting drugs. Interestingly, these 16 codewords in Module B were activated in both immunoglobulin-and ion channel-interacting drugs but followed inverse usage trends: activation was higher in the right subtree for immunoglobulin-related drugs and higher in the left subtree for ligand-gated ion channel-related drugs. In Module C, a set of 24 underutilized codewords out of 128 was associated with seven transmembrane receptors. Other domains also exhibited distinct activation patterns, although the characteristic codewords in these cases are more sparsely distributed.

**Figure 5:**
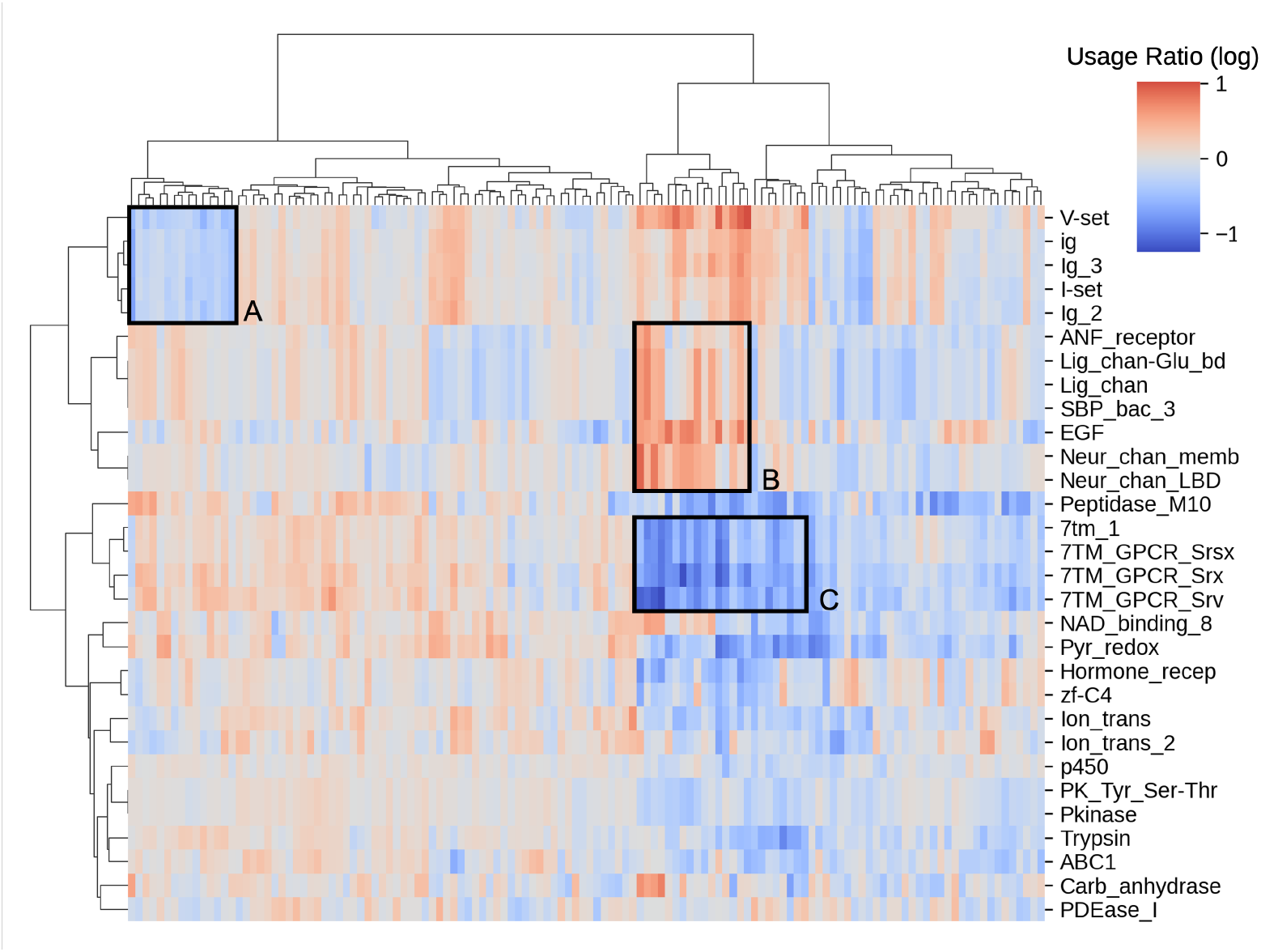
Heatmap showing the usage ratios of drug codewords with respect to interactions with specific protein domains. Each column represents a codeword, and each row corresponds to a protein domain. Boxes indicate characteristic modules identified via hierarchical clustering. Clustered modules are noted as A, B, and C, respectively characterizing immunoglobulin domains: *V-set, ig* (immunoglobulin), *ig 3, I-set*, and *ig 2* ; ligand-gated channels: *ANF receptor, Lig chan-Glu bd* (Ligated ion channel L-glutamate- and glycine-binding site), *Lig chan* (Ligand-gated ion channel), *Neur chan memb* (Neurotransmitter-gated ion-channel transmembrane region), and *Neur chan LBD* (Neurotransmitter-gated ion-channel ligand binding domain); as well as 7 transmembrane receptors: *7tm 1* (7 transmembrane receptor, rhodopsin family), *7TM GPCR Srsx* (Serpentine type 7TM GPCR chemoreceptor Srsx), *7TM GPCR Srx* (Serpentine type 7TM GPCR chemoreceptor Srx), and *7TM GPCR Srv* (Serpentine type 7TM GPCR chemoreceptor Srv).

Taken together, these findings indicate that the codebook, trained through supervised learning, effectively capture domain-level interaction patterns between drugs and proteins. More importantly, even in the absence of additional linear or non-linear transformations, the raw codeword usage itself encodes informative and interpretable signals that are critical for accurate drug–protein interaction prediction.

### 2.6 Bag-of-words(BoW) of codebook captures drug embedding structure without explicit feature semantics

As the codebook stores DTI information, we next aimed to evaluate how well the codeword usage preserves information from the original drug embeddings. In the VQ module, each input representation is divided into multiple slices, with each slice assigned to one of the 128 codewords. This transformation yields a 128-dimensional frequency vector for each drug, where each dimension corresponds to the usage count of a particular codeword. Importantly, this representation discards complex spatial or positional relationships among slices, such as the order or adjacency between codewords, reducing the embedding to a simple frequency profile. This approach is also known as the bag-of-words (BoW) approach [24].

To assess whether the codeword frequency vector could serve as an effective alternative representation of drug features, we compared it with the original drug embeddings using t-SNE for dimensionality reduction and visualization. Specifically, both the original embeddings and the frequency vectors were projected into a two-dimensional space using t-SNE, allowing us to visually inspect the clustering patterns that emerged from each representation. To evaluate the representational capacity of this feature, we annotated the resulting clusters using drug superfamilies as reference categories. As shown in Figure 6, both representations formed clusters that correspond to certain drug superclasses, indicating that meaningful structural or functional groupings were retained even after dimensionality reduction. Notably, the structure derived from the frequency vectors partially resembles those from the original embeddings. This suggests that the frequency-based representation, despite being a simplified and lower-capacity encoding, preserves a substantial amount of the structural information present in the original high-dimensional.

**Figure 6:**
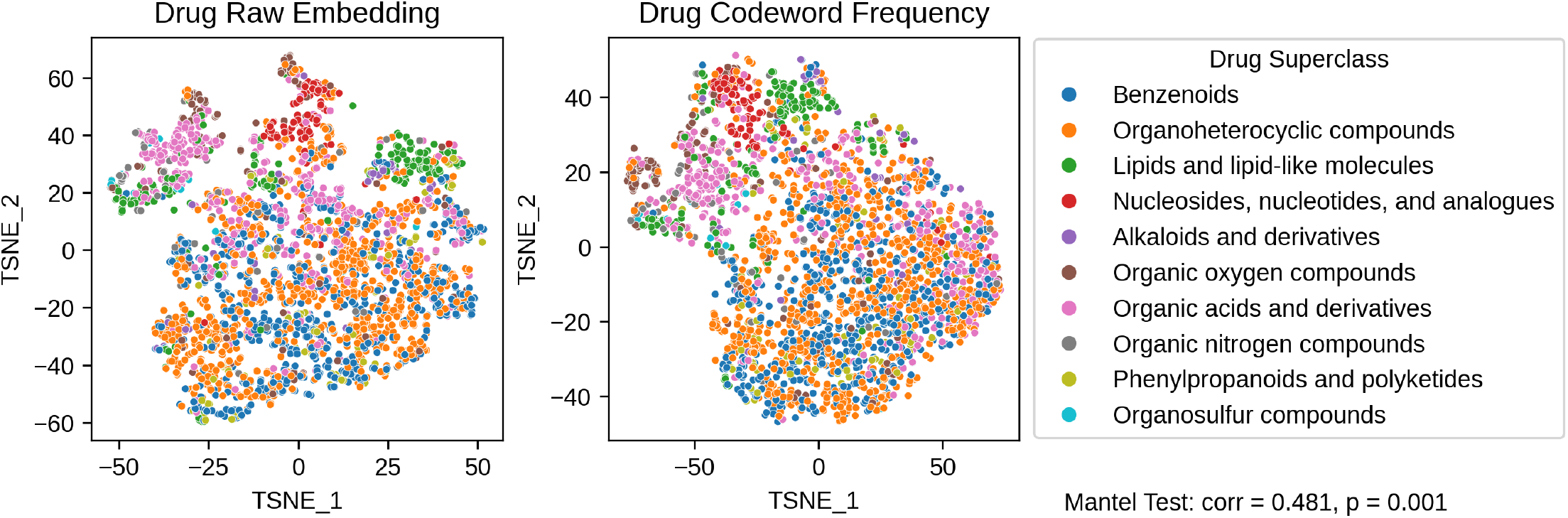
t-SNE projections of drug embeddings generated by the pretrained model (left) and the BoW representation (right). In both plots, drugs are color-coded according to their superclass categories, as curated from the DrugBank database [25].

To quantify this resemblance, we performed a Mantel test to assess the correlation between the pairwise distance matrices of the two representations. The test yielded a correlation coefficient of 0.481 with a highly significant p-value (p = 0.001). It is important to note that accurately grouping drugs by superfamily does not directly reflect the quality of the learned representation, as these categories are not explicitly optimized for in our SVQ framework. Nevertheless, the significant correlation still indicates that even a representation derived solely from codeword frequency counts can retain a substantial portion of the structural information encoded in the original embeddings.

It is worth pointing out that the BoW representation, or the codeword frequency matrix, does not retain any explicit semantic embeddings from the model-generated embeddings. Given this fact, BoW’s ability to preserve the original information strongly suggests that the precise semantic meanings of individual codewords are not essential for representing drugs. Instead, the co-occurrence patterns of codewords appear to encode the most representational value.

To further test this hypothesis, we implemented a variant of the VQ module with a frozen, randomly initialized codebook. Specifically, we applied QR decomposition for orthogonal initialization, ensuring sufficient diversity among the codewords. These codewords remained fixed throughout training. The rationale was that, if the explicit semantic content of the codewords were crucial, freezing the codebook in this way would result in a failure to train the DTI task. Contrary to this expectation, the model remained trainable and achieved less than a 5% performance loss on two large-scale datasets, BioSNAP and BindingDB (Table 4). This result indicates that the specific embeddings within the codebook are not critical. Instead, the model primarily encodes the interaction-relevant information into the co-occurrence patterns of codewords, highlighting that these relational patterns alone are sufficient to capture essential DTI signals. Such a finding not only reinforces the interpretability of our SVQ framework in DTI prediction but also suggests that emphasizing structured co-occurrence over high-dimensional semantic embeddings may represent a broader paradigm in deep learning characterized by both efficiency and interpretability.

**Table 4:** AUPR score for RF and recent-year methods.

| Model | BIOSNAP | BIOSNAP<br>unseen drug | BIOSNAP<br>unseen protein | BindingDB | DAVIS |
| --- | --- | --- | --- | --- | --- |
| frozenVQ-MLP | 0.924 ± 0.001 | 0.886 ± 0.003 | 0.807 ± 0.021 | 0.635 ± 0.007 | 0.401 ± 0.011 |
| SVQ-MLP | 0.928 ± 0.000 | 0.900 ± 0.002 | 0.835 ± 0.003 | 0.668 ± 0.006 | 0.456 ± 0.011 |
| Performance<br>Loss% | 0.43% | 1.56% | 3.35% | 4.94% | 12.06% |

**Table 5:** Summary of Used Datasets.

| Dataset | Split | Drugs | Targets | Interactions | Positive | Negative |
| --- | --- | --- | --- | --- | --- | --- |
| BioSNAP | Train | 4,400 | 2,176 | 19,238 | 9,670 | 9,568 |
|  | Validation | 1,877 | 1,354 | 2,748 | 1,396 | 1,352 |
|  | Test | 2,962 | 1,811 | 5,497 | 2,770 | 2,727 |
|  | Total | 4,510 | 2,181 | 27,483 | 13,836 | 13,647 |
| BindingDB | Train | 3,814 | 1,026 | 12,668 | 6,334 | 6,334 |
|  | Validation | 1,793 | 812 | 6,644 | 927 | 5,717 |
|  | Test | 3,320 | 981 | 13,288 | 1,905 | 11,383 |
|  | Total | 7,164 | 1,254 | 32,600 | 9,166 | 23,434 |
| DAVIS | Train | 68 | 372 | 2,086 | 1,043 | 1,043 |
|  | Validation | 68 | 378 | 3,006 | 160 | 2,846 |
|  | Test | 68 | 379 | 6,011 | 303 | 5,708 |
|  | Total | 68 | 379 | 11,103 | 1,506 | 9,597 |
| BioSNAP<br>unseen drug | Train | 3,588 | 2,177 | 19,154 | 9,535 | 9,619 |
|  | Validation | 1,776 | 1,347 | 2,736 | 1,383 | 1,353 |
|  | Test | 902 | 1,795 | 5,593 | 2,918 | 2,675 |
|  | Total | 4,510 | 2,181 | 27,483 | 13,836 | 13,647 |
| BioSNAP<br>unseen target | Train | 4,383 | 1,744 | 19,375 | 9,876 | 9,499 |
|  | Validation | 1,896 | 1,201 | 2,768 | 1,382 | 1,386 |
|  | Test | 2,931 | 436 | 5,340 | 2,578 | 2,762 |
|  | Total | 4,510 | 2,181 | 27,483 | 13,836 | 13,647 |

## 3 Discussion

In this study, we systematically evaluated the contribution of pretrained features in DTI prediction. Through RF-based feature selection, we demonstrated that not all features provided by language models are equally informative. Linear probing using highly or weakly informative features revealed the risks of feature redundancy and overfitting when high-dimensional representations are not properly filtered, highlighting feature selection as a viable strategy for improving model performance. However, traditional tree-based feature selection methods have problems when integrated into deep learning framework and are often limited in their ability to estimate superposition effects.

Following the principle of reducing feature redundancy and identifying discriminative features, we introduced a VQ module as an alternative. This is because partitioning the high-dimensional, continuous embedding space into multiple subspaces and replacing them with a limited number of discrete latent codewords from a trainable codebook forces the model to learn the most representative and discriminative quantized features in a task-driven manner. This quantization process implicitly reduces redundancy and enhances the expressiveness of the learned representations. Therefore, we regard the VQ method as a soft feature selection strategy.

Compared to linear probing of RF-based Boruta-selected features, the VQ module equipped model (SVQ) achieves competitive performance, outperforming alternatives on three benchmark dataset. While SVQ demonstrates relative lower performance on unseen drugs or targets, it may reflect a trade-off between discrete regularization and generalization. By mapping continuous embeddings to discrete codewords, the VQ module enhances the robustness for familiar input patterns but may struggle to generalize to novel drugs or targets whose representations fall outside the well-optimized codeword regions. This observation highlights an opportunity for further refinement to better balance information preservation and generalization capacity. Nevertheless, our SVQ framework still substantially outperforms recent state-of-the-art models, showing the effectiveness of its architecture in DTI prediction.

Beyond predictive performance, the VQ module also adds a layer of interpretability to the model. Our analysis of codeword usage patterns revealed distinct and domain-specific activation profiles in drugs targeting particular protein domains, indicating that the codebook captures biologically meaningful interaction information, particularly domain-conditioned interactions. Notably, such information is directly reflected in the frequency measure of codewords, without requiring further decoding by downstream linear layers.

During this analysis, we observed that it is not the semantic content of codewords but rather their co-occurrence patterns, that capture the bulk of drug representation and DTI-related information. To address this, we analyzed the performance of a codeword frequency representation, commonly referred to as BoW approach, which discards all semantic content of codewords. It turns out that, this largely simplified representation still preserved a high degree of consistency with the original embedding distribution. It also showed comparable performance with the original embedding in clustering drug categories, resulting in only a marginal drop in the clustering score. These findings provide the evidence for our observation.

To further probe this hypothesis, we conducted an ablation study in which the VQ codebook was randomly initialized and kept frozen during training, thereby preventing the model from learning any meaningful semantic knowledge within the discrete space. In this setting, the model had no access to task-informed codeword semantics, relying solely on the learned co-occurrence patterns of the fixed codewords. If semantic information were indispensable for DTI prediction, this setting would cause the model fail to do correct prediction. However, the results revealed that the model was still able to train successfully and achieved strong performance, with only a slight decrease compared to the fully trained SVQ model.

Our results thus provide strong evidence that it is the co-occurrence of codeword, rather than semantics of codewords, that primarily encodes DTI information. This observation aligns with findings from other domains, such as natural language processing (NLP) [26], where co-occurrence patterns of token usage often convey more information than the semantics of isolated token.

In the above analysis, we focused on the simplest case—a BoW representation. However, additional aspects of codeword interactions remain to be explored, including positional and adjacency relationships between codewords, as well as higher-order interactions across multiple codeword groups. Investigating these richer representational mechanisms represents an important direction for future research.

Overall, our study highlights the importance of selective representation in DTI modeling. We proposed the SVQ framework incorporating a VQ module as a means of approximating feature selection, yielding a model that is both high-performing and interpretable. Notably, the VQ module can be used as an instant plugin on any input embedding pipeline, demonstrating its versatility and broad applicability. As large pretrained language models are increasingly adopted to generate task-agnostic embedding across diverse bioinformatics applications, our findings offer a practical and generalizable strategy for extracting informative features and enhancing model performance.

## 4 Experimental Section

### 4.1 Representation

Two pretrained large language models were used to generate representations of drug molecules and target proteins in this study. Specifically, the Multi-view Molecular Embedding with Late Fusion model (MMELON) [27] was used for drug molecules, while Evolutionary Scale Modeling Cambrian (ESM-C) was applied to target proteins [28].

MMELON is a multi-view foundation model that integrates molecular views of graph, image and text modalities. Each view was pretrained using self-supervised learning on a large scale dataset of 200 million molecules from PubChem and ZINC22, and subsequently combined to enable robust feature learning. During inference, the model takes graph, image and text views, which are derived from SMILES, to produce a 512-dimensional drug embedding, denoted as *E*_*drug*_. MMELON was validated on several protein interaction-related tasks such as human beta-secretase 1 inhibition and cytochrome P450 (CYP) isoform inhibition, where it demonstrated superior robustness in performance, highlighting its potential for broad applications in DTI prediction.

For target proteins, we employ ESM-C, a protein language model optimized for representation learning, and pretrained using a self-supervised masked language modeling approach. To obtain a fixed-length representation, we computed the protein embedding as 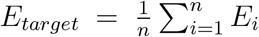, where *n* is the sequence length and *E*_*i*_ is the embedding of the *i*-th amino acid. According to its official benchmark, ESM-C exhibits superior scaling efficiency and outperforms its predecessors with significantly fewer parameters. In our study, we utilized the largest publicly available pretrained ESM-C model, consisting of 600 million parameters, to generate a 1,152-dimensional embedding for each target protein.

Both representations were z-score normalized on a per-dimension basis. Both MMELON and ESM-C model were used in inference mode, and the generated drug and target protein representations were kept frozen during training and testing.

### 4.2 Feature Selection Research

#### 4.2.1 Random Forest Classifier

To conduct the feature selection research, we employed a Random Forest classifier to explore feature contribution of drug and target protein representations for the DTI task. The drug and target protein representation were concatenated to form a unified feature space, where the classifier with 100 decision trees was trained. The Gini impurity was used as the metric to evaluate the importance of representation features.

#### 4.2.2 Boruta Algorithm

After obtaining the feature importance measures, the Boruta algorithm [18] was further applied to determine whether a feature was discriminative or not. This method generates shuffled duplicates of the original features, which are mixed together with the original ones during Random Forest training. Their importance metrics in terms of Gini impurity are compared in each iteration, as an original feature must outperform all other shadow features to be preserved for the next round. This iterative procedure continues until convergence or until a predefined maximum number of iteration is reached, ensuring robust feature selection. In our study, the maximum iteration number was set to 10 to balance computational efficiency with the need for feature evaluation.

#### 4.2.3 Linear probing

After the Boruta iterations, irrelevant features were discarded while discriminative features were retained. Then, as a practice of linear probing research, a simple linear classifier was employed to measure the gain/loss of the representational quality for these discarded or retained features.

The linear classifier was implemented as a multilayer perceptron (MLP) with a linear-Softsign-Dropout-linear structure. The first linear layer projected the concatenated 512-dimensional drug and 1,152-dimensional protein representations into a 512-dimensional latent space, followed by a SoftSign activation function. A dropout rate was set to 0.4. The last linear layer then mapped the features into a 2-dimensional output space, representing class probabilities after a log-softmax transformation. The linear classifier was trained using Negative Log-Likelihood (NLL) loss function.

The model was trained with three different features sets: the full set of features, the Boruta-selected features, and the Boruta-discarded features. To make this comparable, undesired features were masked with zeros to block the gradient from being updated during training. Each configuration was run five times with distinct random initializations. AUPR was used as the primary metric to evaluate representational quality since the benchmark dataset was highly imbalanced.

Our framework incorporated the pretrained large language models, and integrated a VQ module to learn the discrete space of drugs and proteins from their interaction data in a supervised manner.

In the VQ layer, drug and protein embeddings were processed through two separate vector quantizers, where continuous representations were mapped to a fixed number of discrete spaces. Supervision was provided by a fully connected classifier that guided the learning of DTI information, optimizing the capture of relevant patterns in the discrete representations for improved performance on DTI prediction tasks.

More specifically, we employed a split quantization approach [29] for the vector quantizers. Given a continuous embedding *z* ∈ ℝ^*d*^ derived from the drug or protein representation, we had it divided and split into *M* sub-vectors *z*_1_, *z*_2_, … , *z*_*M*_ . Each split *z*_*i*_ with a dimensionality of 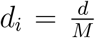 could be mapped to a nearest codeword *e*_*j*_ from a shared codebook

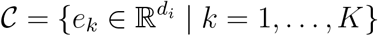

through the VQ approach, which could be defined as

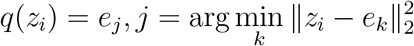

where *q*(*z*_*i*_) is the quantized representation of *z*_*i*_.

The final quantized representation *q*(*z*) was obtained by concatenating the quantized sub-vectors:

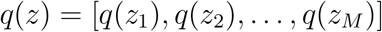

and was passed through the fully connected classifier, which consisted of one linear layer with a Softsign action and another followed by a log-softmax activation. We set hyperparameter *K* = 128 for both drug and protein vector quantizers. We split the drug and protein representations by *n*_*drug*_ = 128 and *n*_*protein*_ = 288, resulting in sub-vector dimensionality equaling 4.

### 4.3 Model Training

Our SVQ training involved the update of the codebooks and the classifier parameters. We designed a hybrid loss function consisting of two components, the codebook loss and the classification loss for the optimizing goal. Firstly, the codebook loss encouraged the quantized representation getting close to the corresponding sub-vectors. Similar to the existing method [30], the codebook loss was defined as follows:

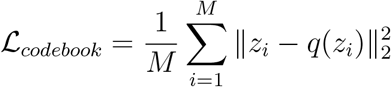

Then, we incorporated the classification loss function to enable drug-target label-guided learning. Given the 2-dimensional output *x* from log-softmax activation, we calculated the loss as NLL loss:

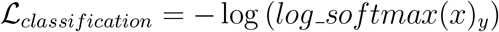

Finally, the loss function was calculated by

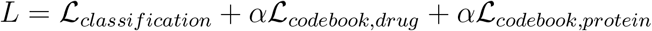

where *α* was the weight of codebook loss set to 0.1.

We trained the model using Adam optimizer with 1e-5 weight decay for 500 epochs. The learning rate was initially set to 0.001 and was adjusted using a cosine annealing schedule, with a restart at the 100th epoch. The learning rate was never reduced below 1e-5. The batch size was set to 128 for all datasets.

### 4.4 Dataset

For model evaluation, we used three widely-used benchmark datasets as summarized in Table 1, including BioSNAP ChG-Miner [31], BindingDB [32], and DAVIS [33]. The structured data setting of these datasets were adopted from ConPLex [12], where positive drug-target pairs were randomly assigned to train, validation and test set with a 7:1:2 ratio.

As described in ConPLex, BioSNAP ChG-Miner contains only known positive drug-target interactions. Therefore, in BioSNAP and its two variants, negative samples were generated by randomly pairing drugs and targets that were not known to interact, under the assumption that such random pairs are highly unlikely to have true interactions. These negative samples were sampled to match the number of positive samples. For BindingDB and DAVIS, drug-target pairs with dissociation constant *K*_*D*_ *<* 30 were labeled as positive, while the rest negative. Notably, these two datasets had balanced train set, but their validation and test sets showed significant positive-to-negative ratio imbalance.

### 4.5 Evaluation Metrics

For statistical comparison of model performance metrics, we employed Student’s t-test, with a significance threshold of *p* < 0.05. To assess the consistency between Bag-of-words representation and the original model-generated embedding, we used the Mantel test, which measured the correlation between two proximity matrices.

## Data Availability

The source code for SVQ is accessible at https://github.com/jdcc2098/SVQDTI.

## Supporting Information

Supporting Information is available from the Wiley Online Library or from the author.

## Author Contribution Statement

J.C. and Y.Z. were responsible for the conceptualization, curation, analysis and visualization of data and writing the original manuscript draft. L.L. and M.W. contributed to the data curation, analysis, and manuscript editing. C.Z. contributed to the data curation and analysis. S.I. and C.L. managed funding acquisition, project administration, supervision, and manuscript editing. All authors approved the final version for submission.

## Acknowledgments

The authors thank the anonymous reviewers for their valuable suggestions. We would like to thank the technical support by the Core Facilities, Zhejiang University School of Medicine. We are grateful to Dr. Hangjun Wu in the Center of Cryo-Electron Microscopy (CCEM), Zhejiang University for his technical assistance on computer clustering.

